# Characterising the Diffusion of Biological Nanoparticles on Fluid and Elastic Membranes

**DOI:** 10.1101/2020.05.01.071761

**Authors:** V.E. Debets, L.M.C. Janssen, A. Šarić

## Abstract

Tracing the motion of macromolecules, viruses, and nanoparticles adsorbed onto cell membranes is currently the most direct way of probing the complex dynamic interactions behind vital biological processes, including cell signalling, trafficking, and viral infection. The resulting trajectories are usually consistent with some type of anomalous diffusion, but the molecular origins behind the observed anomalous behaviour are usually not obvious. Here we use coarse-grained molecular dynamics simulations to help identify the physical mechanisms that can give rise to experimentally observed trajectories of nanoscopic objects moving on biological membranes. We find that diffusion on membranes of high fluidities typically results in normal diffusion of the adsorbed nanoparticle, irrespective of the concentration of receptors, receptor clustering, or multivalent interactions between the particle and membrane receptors. Gel-like membranes on the other hand result in anomalous diffusion of the particle, which becomes more pronounced at higher receptor concentrations. This anomalous diffusion is characterised by local particle trapping in the regions of high receptor concentrations and fast hopping between such regions. The normal diffusion is recovered in the limit where the gel membrane is saturated with receptors. We conclude that hindered receptor diffusivity can be a common reason behind the observed anomalous diffusion of viruses, vesicles, and nanoparticles adsorbed on cell and model membranes. Our results enable direct comparison with experiments and offer a new route for interpreting motility experiments on cell membranes.

## Introduction

Characterising dynamic interactions between nanoobjects and cell membranes is crucial for the understanding of biological processes, such as cell trafficking and viral infection, as well as for the development of therapeutic nanomaterials. The advancements in microscopy techniques, such as single-particle tracking, provide us with the insight into the dynamics of nanooobjects interacting with cell membranes with unprecedented precision. Such studies typically report translational or rotational trajectories of nanoobjects bound to membranes via ligand-receptor interactions. The resulting trajectories can be rarely characterised as random walks, but are typically consistent with some type of anomalous diffusion in which the diffusion coefficient is time-dependent instead of constant [1–9]. This information can in principle serve as a readout of the molecular interactions between the object and the membrane.

Deducing the underlying molecular mechanisms from the trajectories is highly non-trivial as a large number of possible molecular mechanisms can result in similar anomalous motility [10, 11]. For instance, high-speed single-particle tracking studies have reported anomalous diffusion of functionalised nanoparticles, vesicles, and virus-like particles bound to receptors on membranes in living cells [1] and supported bilayers [2–9]. Various physical and chemical effects have been proposed to underlie the observed anomalous diffusion [12], including multivalent interactions between the nanoparticle and receptors, coupling between membrane leaflets [2], molecular pinning [2], receptor clustering [13], formation of transient membrane domains [14–16], and membrane-cytoskeleton interactions [17].

Here we take a reverse approach: we simulate physical interactions between a nanoobject and deformable fluctuating membranes and measure the resulting trajectories for membranes of various properties. We then characterise the resulting diffusion profiles and match them to the underlying molecular mechanisms that are evident in molecular simulations. It is our hope that such an approach can help to interpret experimental trajectories and identify the molecular mechanisms behind them.

We specifically focus on the role of the receptor concentration and the membrane structure/phase state in the resulting diffusion behaviour of the membrane-bound nanoparticle. In our simulations the nanoparticle can for instance represent a virus-like particle, a globular macromolecule, or an inorganic nanoparticle. The particle binds to the membrane via multivalent interactions with the membrane receptors and locally deforms the membrane underneath it. We measure its diffusion profile within molecular dynamics (MD) simulations on fluid, gel-like, and fully cross-linked elastic membranes, at varying receptor concentrations. We find a range of behaviours, from standard random walk to anomalous diffusion characterised by the particle’s hopping between regions of local trapping. We find that the anomalous diffusion is not caused by mutivalent binding, as previously proposed, but by the hindered receptor diffusivity, which results in the particle trapping in regions rich in receptors. The random walk is then recovered if the membrane is fully saturated with receptors. We provide an in-depth numerical analysis of the data and a theoretical framework that characterises the observed anomalous diffusion.

## Methods

This simulation model is schematically depicted in fig. 1. The free-standing membrane is modelled using a coarse-grained solvent-free model, which describes the membrane as a one-particle thick layer of spherical particles (representing a patch of ∼ 10 lipids). To take into account the variety of cell membranes observed in biological systems we employ both a fluid-like membrane model introduced in [18, 19] and an elastic, cross-linked model based on [20]. Values of the parameters are chosen to ensure biologically relevant bending rigidities of ∼ 20*k*_*B*_*T* [18], where *k*_*B*_*T* is the thermal energy. Depending on the choice of the interactions between the membrane-beads, the fluid-like membrane model can capture membranes of different fluidities: from fluid ones in which particles are able to freely diffuse through the membrane to a more gel-like state where diffusion of particles through the membrane is severely limited (see appendix A for specific details of the membrane models).

**FIG. 1:**
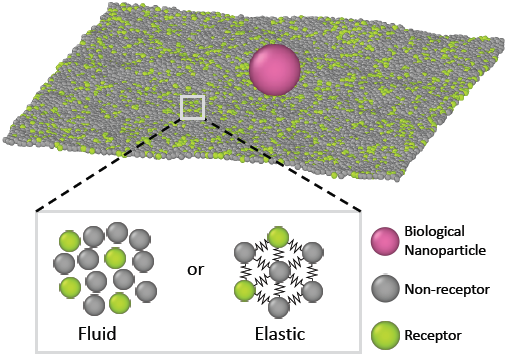
Schematic representation of the system under study. A biological nanoparticle (colored in magenta) adsorbs onto a membrane that is either fluid or cross-linked (elastic). The membrane is described using one-particle thick models, where a certain portion of membrane beads are deemed as receptors (green) and can bind the nanoparticle.

The nanoparticle is represented as a spherical particle with a diameter of 10*σ*, where *σ* is the MD unit of length. It adsorbs onto the membrane and, due to interactions with the membrane particles and thermal fluctuations, diffuses laterally over it. The adsorption is facilitated by allowing certain particles within the membrane to bind the particle. These are deemed as receptors and attract the particle via a truncated and shifted Morse potential. Both receptors and non-receptors (non-binding membrane particles) interact with the nanoparticle via volume exclusion (see appendix B for more details).

### Simulation Details

Each simulation starts by placing 7020 (fluid) or 7832 (elastic) membrane particles on a planar hexagonal lattice, randomly labeling a chosen percentage of them as receptors. The nanoparticle is placed above the center of the membrane and numerical integration is carried out with Langevin dynamics [21] within the *NPH* (constant particle number, pressure, and enthalpy) ensemble in LAMMPS [22]. Periodic boundary conditions are imposed and every run consists of at least 10^7^ iterations with a timestep Δ*t* = 0.01*τ* (fluid) or Δ*t* = 0.009*τ* (elastic) where *τ* denotes the time unit of the system. The first 5 · 10^5^ iterations serve to equilibrate the system after which the time origin is set and the lateral position of the nanoparticle **r**(*t*_*n*_) can be tracked at discrete times *t*_*n*_ = *n*Δ*t* with *n* being the number of timesteps.

### Diffusion on Fluid Membranes

Owing to the free diffusion of receptors, the fluid membrane will encapsulate the nanoparticle when it consists of too many receptors (≳ 40%), which severely hinders its lateral motion (see supplementary movie 1). To explore the motion of the nanoparticle before the encapsulation, we chose to fix the receptor percentage at 20%. For this receptor percentage we observe clear lateral trajectories of the nanoparticle (see fig. 2 and supplementary movies 2 and 3), where it is readily seen that the trajectories on a fluid membrane are significantly different from the ones on a gel-like membrane. On the gel-like membrane the motion appears to be more clustered and alternating between parts of confined motion and rapid hops in between, effectively dividing the trajectories in mobile and immobile segments. This becomes especially clear by noting the plateaus in the 2D radial distance *r* = |**r**(*t*) − **r**(0) | over time (with **r**(*t*) being the lateral position of the nanoparticle at time *t*). Motion on the fluid membrane seems, in contrast, to be less clustered and more reminiscent of normal diffusive behavior.

**FIG. 2:**
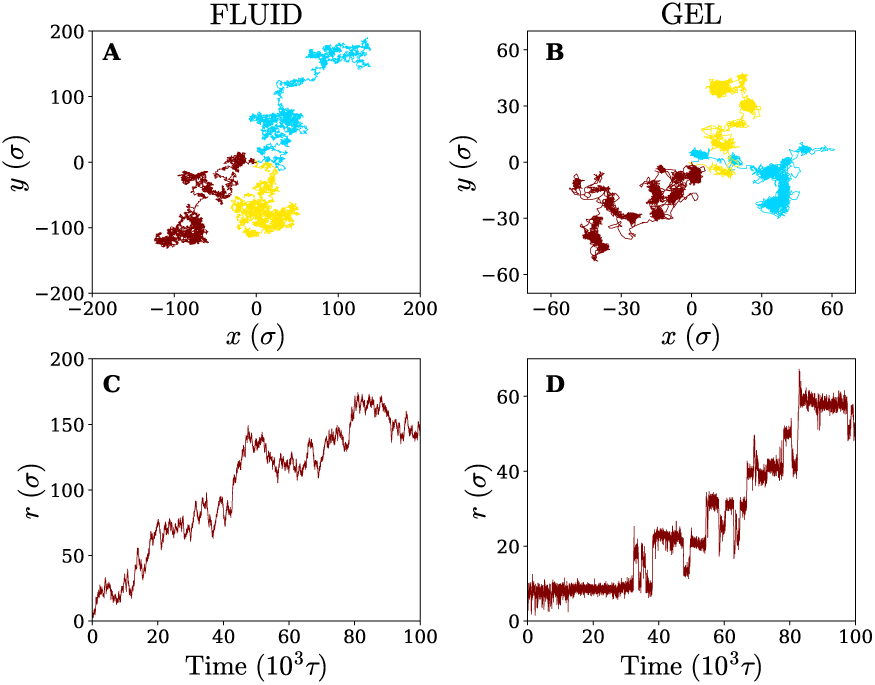
Diffusion on fluid and gel-like membranes. (a,b) Example trajectories of the nanoparticle on (a) fluid and (b) gel-like membranes. (c,d) 2D radial distance *r* as a function of time for the trajectories shown in brown, on (c) fluid and (d) gel-like membranes.

To better quantify the difference between the nanoparticle motion on the fluid and gel-like membrane we have plotted the (time-ensemble averaged) mean square displacement (MSD), i.e. ⟨ (**r**(*t*) − **r**(0))^2^ ⟩, and the corresponding instant diffusion coefficient *D* = MSD*/*4*t* for the particle (fig. 3). It can be noticed that after a brief increase due to ballistic motion, the instant diffusion coefficients *D* start to decrease over multiple orders of magnitude of time. This decrease is almost negligible for the fluid membrane which corresponds to normal diffusion, where MSD ∝ *t*, and thus a constant *D* [23]. For the gel-like membrane the diffusion coefficient is qualitatively different since it drops approximately an order of magnitude before saturating towards a constant value. In other words, clear anomalous diffusion occurs on the observed time scales, while normal diffusion is regained in the long time limit. Interestingly, similar behavior has been observed in experiments involving the diffusive motion of a spherical gold nanoparticle binding to a model membrane via receptor-ligand interactions (see fig. 3). It thus seems that the temporal trapping of the nanoparticle, sometimes of the order of the simulation time (explaining the relatively large variety between time averaged MSDs), results in anomalous diffusion on membranes of decreased fluidities.

**FIG. 3:**
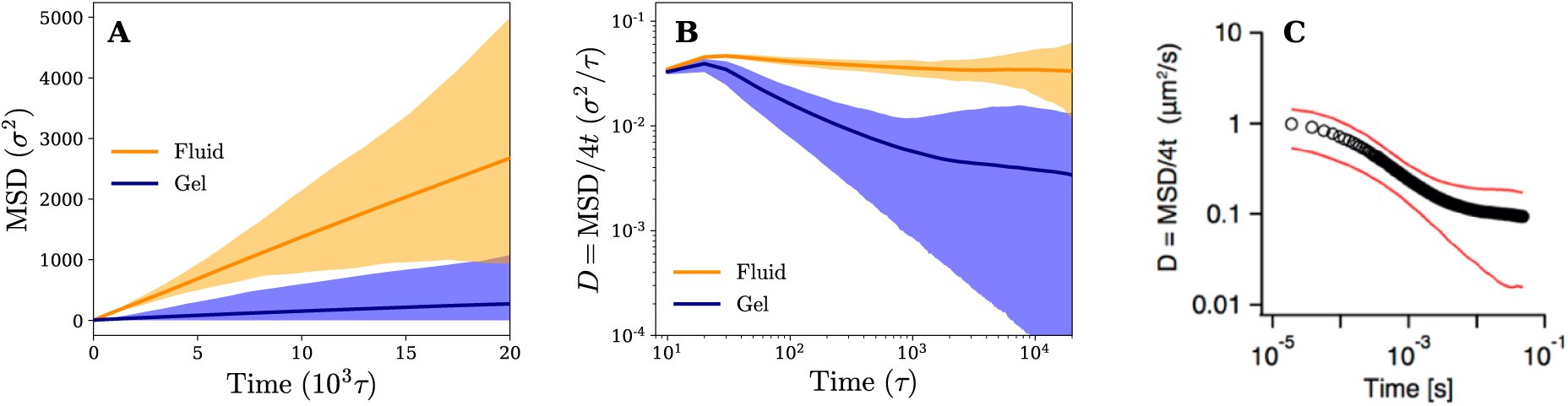
Diffusion on fluid and gel membranes is qualitatively different. (a) Time-ensemble averaged MSD in the lateral direction for diffusion on a fluid (in orange) and gel-like membrane (in blue). (b) The instant diffusion coefficient derived from the time-ensemble averaged MSD from (a). The shaded areas represent the variation in time averaged MSDs and corresponding instant diffusion coefficients. (c) The experimental result of a 40 nm gold particle that binds to a supported model membrane. Reprinted with permission from [2]. Copyright 2014 American Chemical Society.

The analysis of the MSD is, however, limited and can only identify whether the motion is anomalous or not. It gives little information about the underlying mechanism of the observed diffusion process. To further rationalise our findings, we have therefore retrieved the (2D) cumulative density function (CDF) *P* (*r, t*), i.e. the probability that a particle starting at the origin is found within a circle of radius *r* at time *t*, for several discrete times *t*_*n*_ = *n*Δ*t*. This can be achieved by counting the number of absolute displacements |**r**(*t*_*i*+*n*_) − **r**(*t*_*i*_) | < *r* for all trajectories and normalising over the total number of considered data points [24]. The results for *t*_*n*_ = 1000*τ* are shown in fig. 4, which again demonstrate qualitatively different behavior for the fluid and the gel-like membrane settings.

**FIG. 4:**
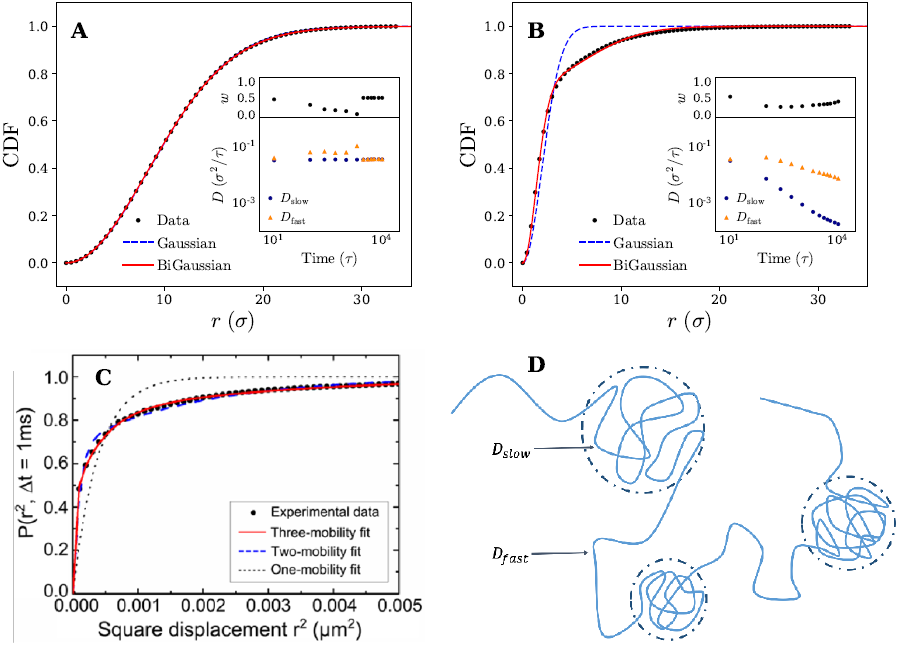
Diffusive modes on fluid and gel-like membranes. Plots of the cumulative density function *P* (*r, t*) for (a) fluid and (b) gel membranes. Fits to both the Gaussian (eq. (1)) and bi-Gaussian (eq. (2)) result are plotted for *t*_*n*_ = 1000*τ*. Insets show *D*_fast_, *D*_slow_, and *w* as a function of time resulting from a bi-Gaussian fit of the retrieved CDFs. (c) Experimental results of the cumulative density function (CDF) along with fits to the one-mobility (Gaussian), two-mobility (bi-Gaussian) and three-mobility models for the diffusion of a 20nm gold nanoparticle attached to GM1 ganglioside in supported DOPC bilayer. Reprinted with permission from [13]. Copyright 2014 American Chemical Society. (d) Schematic representation an example nanoparticle trajectory indicating how it alternates between temporary confined parts and fast diffusive parts. These different segments of the trajectory can be characterised by a slow and fast diffusion coefficient respectively (*D*_slow_ and *D*_fast_) which have also been shown in the picture.

We assess this discrepancy more quantitatively by introducing a two-component mobility model [24–27] in which the obtained CDFs are fitted with both a single Gaussian distribution corresponding to normal diffusion (Brownian motion)

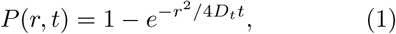

and a bi-Gaussian distribution given by

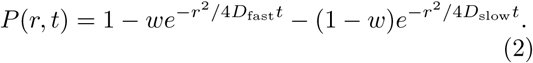

The model thus allows us to identify if a single, ‘simple’ diffusive process with a diffusion coefficient *D*_*t*_ is sufficient to describe the (anomalous) motion or that (at least) two distinct processes, characterised by a slow *D*_slow_ and fast *D*_fast_ diffusion coefficient with a relative weight *w*, are required.

Fits to eqs. (1) and (2) are shown in fig. 4 and indicate that for the fluid membrane settings both distributions provide high quality fits, confirming that particle motion on the fluid membrane is governed by normal diffusion. However, on the gel-like membrane only the bi-Gaussian is able to accurately fit the observed results, which is again in qualitative agreement with experimental results of a gold nanoparticle diffusing on a model membrane, as shown in fig. 4. The particle motion is thus effectively split in two (or possibly more) distinct diffusive modes which, combined, are likely responsible for the anomalous diffusion manifested in the MSD.

To gain more insights from the fits and study the observed behavior over time, we have plotted the bi-Gaussian fitting parameters (*D*_fast_, *D*_slow_, and *w*) over a range of times (see insets fig. 4). It can be seen that on the fluid membrane, as expected, normal nanoparticle diffusion occurs on all considered timescales. In particular, the fast and slow diffusion coefficient remain either equal to each other, which is the same as a single Gaussian fit, or differ only slightly.

Switching to the gel-membrane settings, we find that in this case normal diffusion (approximately equal *D*_slow_ and *D*_fast_) is only obtained for the smallest considered time (*t*_*n*_ = 10*τ*). Careful inspection of the trajectories (fig. 2) suggests that this is a consequence of the nanoparticle not yet experiencing the effects of the temporal confinements. At later times, however, the values of the slow and fast diffusion coefficient become increasingly separated and a clear distinction can be made between both diffusive modes. Interestingly, both diffusion coefficients are decreasing, but the slow diffusion coefficient drops much faster in value (over two orders of magnitude). A possible explanation for this rapid drop in *D*_slow_ and the slower decrease in *D*_fast_ can be found when we link the slow diffusion coefficient to the confined parts of the trajectories (see fig. 4 for a schematic representation). Since the MSD saturates for confined diffusion [28], the slow diffusion coefficient is expected to decay to zero in the long time limit, i.e. when the time is much larger than the average confinement time. Consequently, the overall diffusive motion of the nanoparticle should tend (in the same limit) towards normal diffusion with a smaller diffusion coefficient, which explains the more slowly decreasing (and possibly saturating) *D*_fast_.

We finalise our analysis of the nanoparticle motion and corresponding underlying diffusion process by calculating the normalised velocity auto-correlation function (VAF). This quantity, often used as a diagnostic tool to distinguish among different mechanisms for anomalous diffusion, is defined as [29–31]

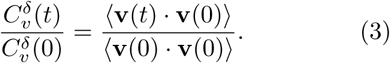

Here brackets denote time-ensemble averaging and the velocity of the nanoparticle at time *t* is calculated over a time period *δ* via

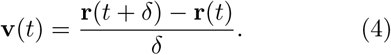

The results for different times *δ* have been plotted in fig. 5. On the gel-like membrane we notice the formation of clear negative peaks in the VAFs at *t* = *δ* over a long range of *δ* values. This indicates the existence of antipersistent behavior, which is likely a result of the temporary confinement experienced by the nanoparticle. In particular, the tendency of the particle to move back into its temporary confined area and visit places it has visited before, negatively correlates the velocity.

**FIG. 5:**
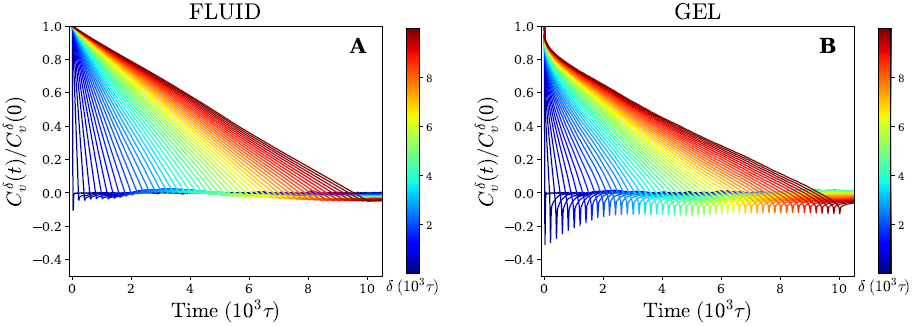
Antipersistent motion on a gel-like membrane. Velocity autocorrelation function for (a) fluid and (b) gel membrane as defined by eq. (3) (using time-ensemble averaging) for different times *δ* that indicate the time over which the velocity is defined (eq. (4)).

For the fluid membrane settings we cannot identify significant antipersistent behavior and the VAFs decay to zero at *t* = *δ*. This is consistent with normal diffusion for which the direction of movement is random and no correlations between the velocity at different times are expected.

### Diffusion on Elastic Membranes

Having observed that anomalous diffusion occurs more prominently upon making a fluid model membrane more gel-like, we now study the nanoparticle motion in the limit of a completely cross-linked membrane, where membrane components cannot diffuse even at infinitely long times. Such membranes are a good model for the membrane of red blood cells, which contain actin-spectrin networks [32], or for thin elastic shells and thin polymer films [33–36].

In comparison to its fluid counterpart, the elastic membrane, due to its cross-linked nature, never encapsulates the particle, even when it consists solely of receptors. This allows us to study the diffusion of the particle across the membrane for different receptor percentages. Letting the trajectories again serve as our starting point (see fig. 6), we find that for 30% receptors (smaller percentages lead to particle detachment) clear clustering and temporal confinement of the particle occur; these hallmarks become even more evident at a larger receptor percentage of 50% (increased membrane inhomogeneity). However, on a homogeneous membrane consisting of only receptors the trapping behavior disappears. The particle motion on the elastic membrane therefore seems, despite the change in membrane mechanics, to be very similar to that on a fluid (gel-like) membrane. The only difference rests in the fact that, instead of receptor diffusivity, the percentage of receptors appears to control the transition from hop-like diffusion towards a freely diffusive behavior. This clearly indicates that the membrane inhomogeneity, rather than the multivalent binding between the particle and the receptors, is responsible for the observed anomalous behaviour.

**FIG. 6:**
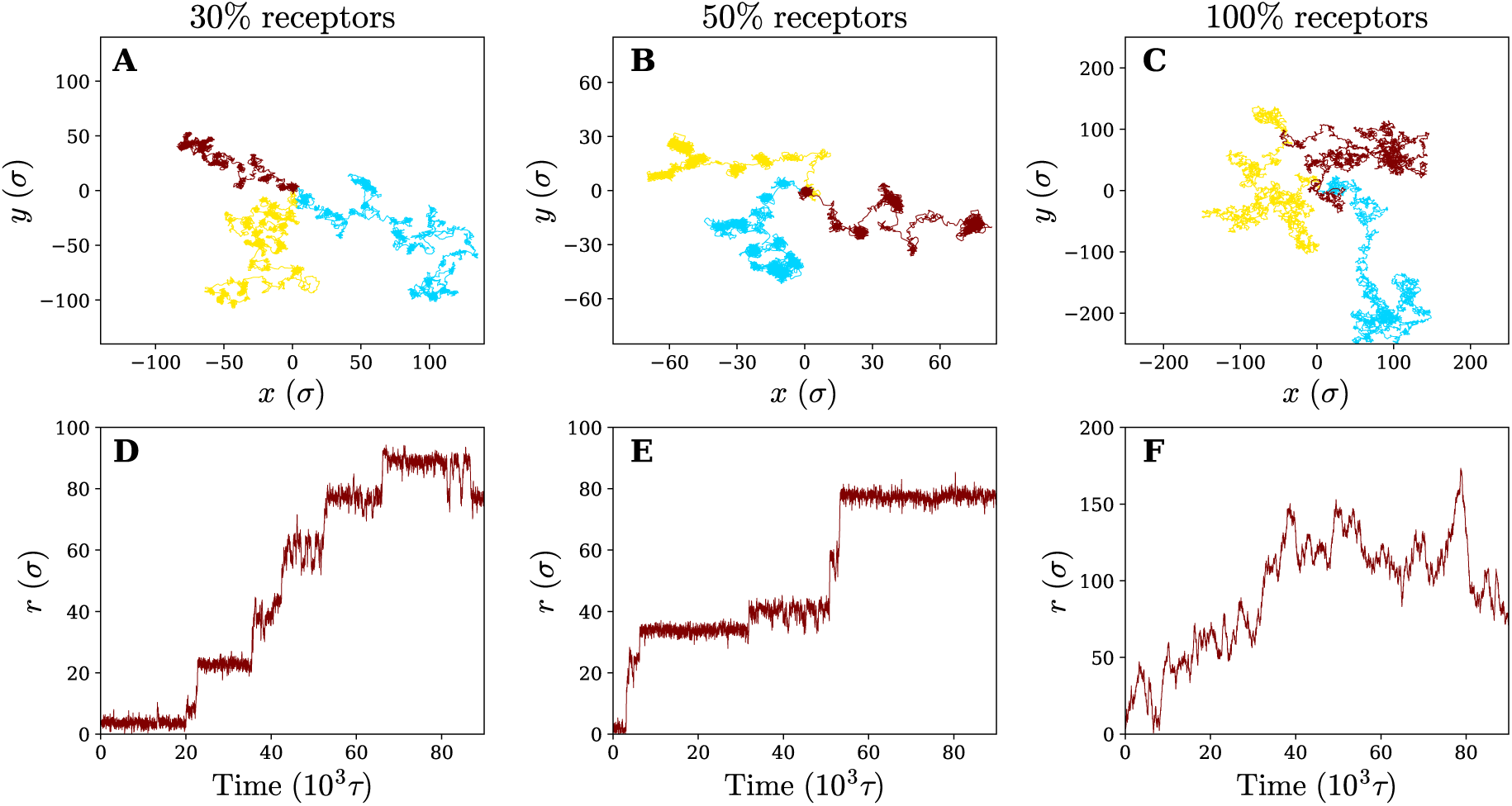
Diffusion on an elastic, cross-linked, membrane. Three example trajectories of the nanoparticle on an elastic membrane of varying receptor percentages: (a) 30% (b) 50 % and (c) 100 %. Corresponding 2D radial distance as a function of time for the green trajectory (d,e,f).

To study the effect of the receptor percentage in more detail and test whether the observed particle motion is truly similar for both membranes, we have calculated the same quantities that have been used to characterise the diffusion process on fluid and gel-like membranes. The (time-ensemble averaged) MSD and related instant diffusion coefficient *D* = MSD*/*4*t* on the elastic membrane are shown in fig. 7. They demonstrate experimentally consistent subdiffusive behavior (decreasing *D*) on intermediate timescales for 30% and 50% percent receptors, as opposed to normal diffusive behavior (constant *D*) for 100% receptors. This again corroborates that the temporal trapping of the particle underlies anomalous diffusion. Since on average the particle trajectories involve more clustering and longer trapping at 50% receptors compared to 30%, this also explains why the drop in value of *D* is larger for the former.

**FIG. 7:**
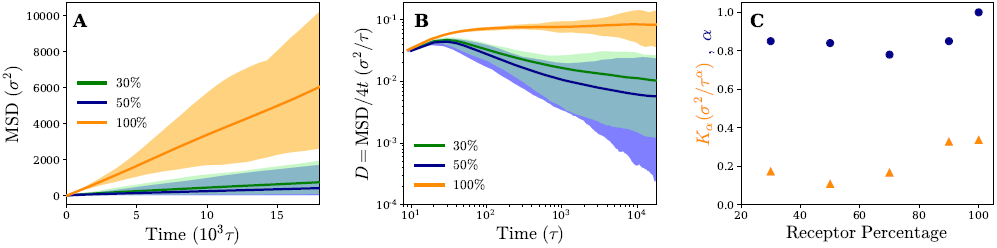
Diffusion is governed by the pattern of receptors, rather than the membrane’s mechanics. (a) Time and time-ensemble averaged MSD in the lateral direction (*x, y*) for the nanoparticle diffusion on an elastic membrane for different percentages of receptors. (b) The instant diffusion coefficient derived from the 2D time-ensemble averaged MSD with the shaded area representing the variation observed in the time averaged MSDs. Results are plotted for different receptor percentages. (c) The anomaly exponent *α* and anomalous diffusion coefficient *K*_*α*_ as a function of receptor percentage. Results are obtained from a least squares fit of the corresponding time-ensemble averaged MSD.

The fact that the subdiffusive behavior initially becomes more apparent with increasing receptor percentage, but diminishes when going to a membrane consisting of only receptors, hints at some non-trivial dependence of the motion on receptor percentage. We quantify this by retrieving the MSD at a number of different receptor percentages and fitting the results to MSD = *K*_*α*_*t*^*α*^. The obtained anomalous diffusion coefficients *K*_*α*_ and anomaly exponents *α* are plotted in fig. 7. It can be seen that, starting from 30% receptors, anomalous behavior and weaker diffusion become more apparent at first (smaller values of *α* and *K*_*α*_), most notably around 50% − 70%, but eventually fade away leading to normal diffusion for 100% receptors. These results suggest that there exists an optimum receptor percentage around ∼ 50% for which anomalous diffusion is most evident in completely cross-linked membranes.

Moving back to the nature of the anomalous particle diffusion, we have examined the CDFs for elastic membranes consisting of 50% and 100% receptors by means of the two-component mobility model (see fig. 8). At an intermediate time of *t*_*n*_ = 900*τ*, we note that both a Gaussian (eq. (1)) and bi-Gaussian (eq. (2)) give identical high quality fits to the CDF on a uniform membrane (100% receptors). The accompanying fit parameters of the bi-Gaussian (*D*_fast_, *D*_slow_, and *w*) in turn show that this is the case across the entire analysed time range, since the fast and slow diffusion coefficient remain (almost) equal to each other. Realising that over time the fitted diffusion coefficients also keep an approximately constant value, which is in quantitative agreement with the one derived from the MSD, we conclude that the motion corresponds to normal diffusion.

**FIG. 8:**
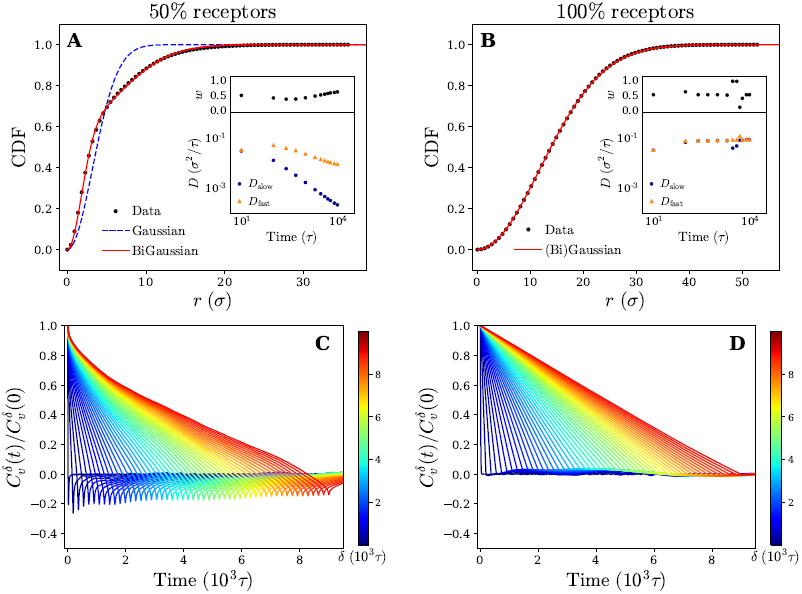
Diffusive modes and antipersistent motion on an elastic membrane. Plots of the cumulative density function *P* (*r, t*) at time *t*_*n*_ = 900*τ* for an elastic membrane that contains (a) 50% and (b) 100% receptors. Fits to both the Gaussian (eq. (1)) and bi-Gaussian (eq. (2)) result are plotted as well. Insets show *D*_fast_, *D*_slow_, and *w* as a function of time resulting from a bi-Gaussian fit of the retrieved CDFs for several times *t*_*n*_. (c) and (d) Plots of the velocity autocorrelation function as defined by eq. (3) (using time-ensemble averaging) for different times *δ* that indicate the time over which the velocity is defined (eq. (4)) for a membrane that contains (c) 50% and (d) 100% receptors.

**FIG. 9:**
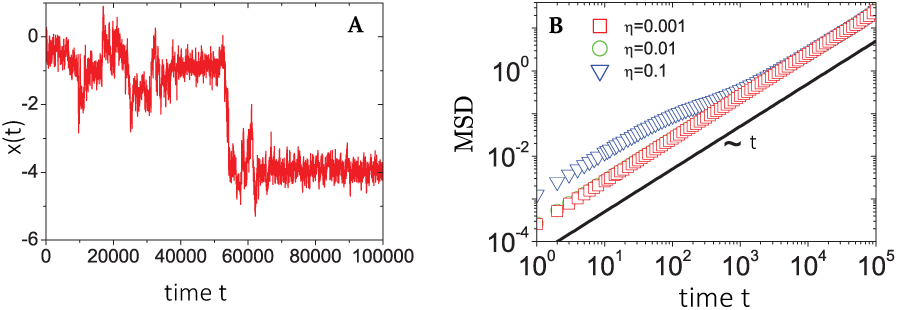
Plots of (a) a sample trajectory 1D *x*(*t*) and (b) the time-ensemble averaged MSD of a nCTRW with Ornstein-Uhlenbeck noise. MSDs are retrieved for increasing noise strengths *η*, while the trajectory corresponds to the largest noise strength *η* = 0.1. Figures are adapted from [42], with the permission of AIP Publishing.

On a non-uniform membrane with 50% receptors we see that at a time of *t*_*n*_ = 900*τ* (within the subdiffusive regime of the MSD) the CDF deviates significantly from a single Gaussian. A bi-Gaussian, in comparison, is able to accurately fit the CDF suggesting that the motion is split in two or possibly more diffusive modes. Moreover, the shape of the CDF curve clearly resembles the one obtained for a gel-like membrane (see figs. 4 and 8). Combined with the already observed similarities between both membranes in terms of trajectories and MSD, we expect the underlying mechanism of the anomalous particle motion on a non-uniform elastic membrane to be the same as on the gel-like membrane. This claim is further substantiated by considering the bi-Gaussian fit parameters (*D*_fast_, *D*_slow_, and *w*), which change over time in an almost identical manner as those for the gel-like membrane (see insets figs. 4 and 8). Applying earlier proposed reasoning, we can again explain the fast drop in *D*_slow_ and more moderate decrease of *D*_fast_ by linking the former to the confined parts of the particle trajectories.

Finally, for completeness, we have also calculated the VAFs corresponding to both a membrane composition of 50% and 100% receptors. The results are plotted in fig. 8 and appear akin to the ones obtained for a gel-like and fluid membrane respectively. Specifically, we see significant negative correlations (antipersistent behavior) for the 50% membrane, which can be attributed to the temporary confinement of the nanoparticle. These results thus provide additional support that the underlying diffusion mechanism of the nanoparticle is independent of our choice of membrane model and rather seems governed by receptor patterning and diffusivity.

### Anomalous Diffusion Models

Since anomalous diffusion, i.e. stochastic motion that differs (significantly) from the laws of ‘simple’ Brownian motion, is being observed in an increasing amount of both animate and inanimate systems, several theoretical models have also emerged in literature to capture the physical origins of the exhibited anomalous motion [37]. Accordingly, we also seek to provide a more general framework for our simulation results by discussing the observed anomalous diffusion in the context of such models, concentrating on three popular examples: fractional Brownian motion (fBM), confined Brownian motion (cBM), and a continuous time random walk (CTRW) [38, 39].

Fractional Brownian motion is often used to describe stochastic motion within a viscoelastic medium. In this case the particle locally deforms the medium, which results in a tendency for it to go back to locations it has visited in the past and in turn yields antipersistent behavior. Although this is consistent with the obtained simulation results, fBM also predicts a single Gaussian cumulative density function (CDF) and a subdiffusive (time-ensemble averaged) MSD in the long time limit [38], which are both incompatible with the obtained results. The second model, cBM, describes normal diffusion within a form of confinement (e.g. hard walls or a harmonic potential). This yields antipersistent behavior, but also a saturating MSD which is not observed in our simulations. The most intuitive mechanism to describe the simulation results with is the CTRW. This model describes the motion of a particle in terms of random jumps Δ**r**, drawn from a distribution *p*(Δ**r**). In between these jumps the particle is immobilised for a certain waiting time *t*_*w*_, which is drawn from an independent distribution *ψ*(*t*_*w*_) [40, 41]. This notion of temporal confinement lines up with our particle trajectories. However, due to the assumed random jump direction in a CTRW, no antipersistent behavior is expected to occur [30] and therefore it disagrees with our simulation results.

It thus seems that the proposed single models are inadequate to rationalise our simulation results. An alternative approach, inspired by the accurate bi-Gaussian fit of the cumulative density functions, is to segment trajectories in distinct diffusive modes and assign separate models to each of them [30], or to combine different anomalous diffusion models [42]. We focus on the latter by invoking a so-called noisy continuous time random walk (nCTRW) [37, 42]. In principle, a nCTRW combines the CTRW and cBM processes by superimposing Ornstein-Uhlenbeck (OU) noise, i.e. diffusion in a harmonic potential, upon the CTRW motion such that the particle jiggles around its CTRW position during the waiting time (see appendix C for more details). Supported by our trajectories, we can relate these two separate mechanisms to the temporal binding and fast jumps of the nanoparticle (CTRW), and the thermal fluctuations it still experiences while being bound (OU noise). In fact, trajectories obtained from simulating a nCTRW appear akin to the ones obtained for our nanoparticle (see appendix C). Besides linking to the observed trajectories, the nCTRW also supports the bi-Gaussian fits of the CDF (two distinct diffusive modes) and explains the observed antipersistent behavior of the VAF since the OU process tends to drive a particle back towards its ‘binding site’ set by the CTRW. Additionally, numerical simulations and an analytical derivation (appendix C) indicate that, for strong enough OU noise, the time-ensemble averaged MSD of a particle subject to a nCTRW increases sublinearly (anomalous diffusion) over an intermediate time range before saturating towards a linear increase in time in the long time limit [37, 42]. This makes our results consistent with the theoretical description of a nCTRW, which thus provides a promising (combined) mechanism to describe the diffusive motion of temporally bound, thermal particles. Overall, we have demonstrated that, already for a coarse-grained simulation set-up, theoretical models require a superposition of stochastic processes to fully grasp the complexity often encountered in biological systems.

## Discussion and Conclusion

We have used coarse-grained molecular dynamics simulations to investigate diffusion of a nanoobject adsorbed onto fluid, gel-like, and cross-linked elastic membranes of varying receptor concentrations. The nanoparticle binds receptors in the membrane via multivalent interactions and captures several tens of receptors at any point. By doing so the particle deforms the membrane underneath it, to a height of ∼ 10% of its diameter. This type of interaction however does not cause anomalous diffusion on fluid membranes, as the bound receptors (and effective membrane deformation) are able to diffuse together with the particle [43]. Interestingly, even at higher receptor concentrations, when the particle becomes well-wrapped by the membrane, to the point of endocytosis, we have found that the diffusion retains its normal profile, albeit at a lower value of a diffusion coefficient (see supplementary movie 1).

In contrast, if the membrane fluidity is decreased such that the receptors cannot freely diffuse anymore, the anomalous diffusion naturally arises. The particle diffuses normally until it hits a region with a higher local receptor concentration where it gets trapped, resulting in a lower local diffusion coefficient. Occasionally, the particle hops between such regions, which gives rise to a second, higher diffusion coefficient, and overall a trajectory that is inconsistent with a random walk. This behaviour is more pronounced at higher receptor concentrations, in good agreement with the experimental study of Hsieh *et al*. [13] for a system of a gold nanoparticle bound to GM1 ganglioside or DOPE lipids in supported DOPC bilayers. Specifically, they also reported trajectories of strong transient confinements and overall anomalous diffusion, which could be decomposed into two effective diffusion coefficients. Furthermore, their study showed that the anomalous diffusion is more pronounced at higher receptor concentrations, consistent with our simulations. Very similar results to ours were also reported in high-speed single-particle tracking of a gold nanoparticle bound to GM1 receptors in model membranes [2].

Our study suggests that the change in the membrane structure, and/or the inability of the receptor to freely diffuse, can possibly explain a plethora of previously reported experimental results and drive new studies. The conclusion can be easily tested in membranes of controlled fluidities [44]. Our approach is different to previous numerical approaches e.g. [10, 12, 17] as it particle-based: it incorporates explicit molecular ingredients, their interactions and mechanics. Our simulation set-up can easily incorporate additional effects regularly found in more biologically-realistic settings, such as heterogeneity in the lipid composition [45], direct interactions between membrane proteins [46], asymmetry of the nanoparticle ligand arrangement [47], or the presence of membrane-deforming machinery [48]. We hope that our study will inspire future feedback between experimental studies of membrane-adhering components and coarse-grained simulations.

## Acknowledgements

We thank Jessica McQuade for her input at the start of the project. We acknowledge support from the ERASMUS Placement Programme (V.E.D.), the UCL Institute for the Physics of Living Systems (V.E.D. and A.Š.), the UCL Global Engagement Fund (L.M.C.J.), and the Royal Society (A.Š.).

## A Membrane Models

Both the fluid and elastic model membrane consist of a one-particle thick layer of spherical particles with diameter *σ* and mass *m* (which, along with the thermal energy *k*_*B*_*T*, set the length, mass, and energy scale of the system). Their different nature arises as a result of distinct particle-particle interactions to form a coherent membrane.

### Fluid Model

In the fluid model the membrane particles interact with one another through a pairwise interparticle potential given by [18, 19]

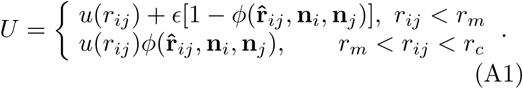

Here **r**_*ij*_ denotes the distance vector between particles *i* and *j* with *r*_*ij*_ = |**r**_*ij*_| and 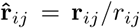, while the unit vectors **n**_*i*_ and **n**_*i*_ represent the axes of symmetry of particle *i* and *j* respectively. We set *ϵ* = 4.34*k*_*B*_*T, r*_*m*_ = 2^1*/*6^*σ*, and *r*_*c*_ = 2.6*σ*. The potential *U* is separated in a distance-dependent part *u*(*r*) that forces particles to stick together and an orientation-dependent part 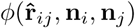 which substitutes the hydrophobic effects of the lipids. The former of these is described by

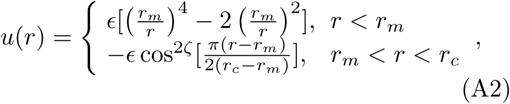

where the repulsive branch (*r* < *r*_*m*_) is given by a ‘soft’ 4-2 Lennard-Jones (LJ) potential. The attractive branch (*r*_*m*_ < *r* < *r*_*c*_) is a cosine function that decays to zero at the cutoff radius *r*_*c*_. The exponent *ζ* determines how rapidly the function tends to zero and serves as a measure of the diffusivity of the membrane particles. Its value is set at either *ζ* = 4.0 to obtain a fluid state where particles can freely diffuse through the membrane, or *ζ* = 2.5 resulting in a more gel-like membrane state where diffusion of particles is severely limited [18, 19]. More precisely, this results in particle diffusion coefficients of the order of ∼ 0.1*σ*^2^*/τ* and ∼ 10^−4^*σ*^2^*/ τ* respectively with *τ* = (*mσ*^2^*/k*_*B*_*T*)^1*/*2^ the time unit of the system [18]. The orientation-dependent function yields

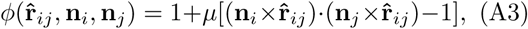

and drives the membrane particles towards a flat configuration w ith i ts d irect s urroundings. Note that this form neglects spontaneous curvature and we thus assume the membrane to be locally flat on the investigated length scales, i.e. only a small part of the cell membrane is modelled. The parameter *µ* = 3.0 can be interpreted as a weight of the energy penalty when the particles are deviating from a flat configuration, and therefore relates to the bending rigidity of the model membrane.

### Elastic Model

The elastic model is created by placing all particles on the nodes of a standard triangulated mesh with hexagonal symmetry [49]. To avoid overlap, each pair of particles repels one another through a Weeks-Chandler-Andersen (WCA) potential, i.e.

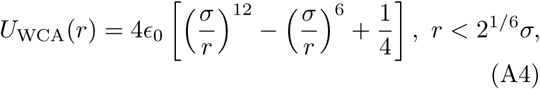

where *ϵ*_0_ = 5.0*k*_*B*_*T* and *r* denotes the distance between the centers of two membrane particles. We then enforce surface fixed connectivity to the membrane by connecting each particle with its six nearest neighbours via a harmonic spring potential (see fig. 1)

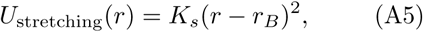

which is described in terms of the spring constant *K*_*s*_ = 18*k*_*B*_*T/σ*^2^ and an equilibrium bond length *r*_*B*_ = 1.23*σ*. These model the stretching rigidity and the equilibrium configuration of the membrane respectively. To ensure an energetically favorable flat membrane, the model includes a bending rigidity in terms of a dihedral potential between adjacent triangles on the mesh

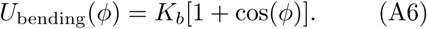

Here *K*_*b*_ = 20*k*_*B*_*T* is the bending constant and *ϕ* the dihedral angle between opposite vertices of any two triangles sharing an edge.

### B Nanoparticle-Membrane Interaction

The employed model membranes are composed of two types of particles, i.e. receptors and non-receptors. To prevent overlap and enforce volume exclusion, both these type of particles interact with the nanoparticle via a scaled WCA potential *U*_WCA_(*R* − Δ) [see eq. (A4)], where *R* denotes the center-to-center distance between the nanoparticle and the respective membrane particle, and Δ = 4.5*σ* the difference between the radii of both particles. Receptors distinguish themselves from non-receptors by also attracting the nanoparticle via a truncated and shifted Morse potential, i.e.

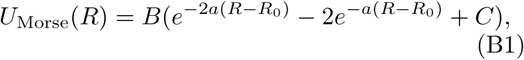

for *R* ≤ 10*σ* and zero otherwise. The attractive strength *B* = 1.5*k*_*B*_*T* and width of the potential *a* = 3.0*σ*^−1^ have been chosen such that in all simulations the nanoparticle stays bound to the membrane (no detachment) but can still laterally move over it (no permanent binding at one site). Moreover, we let the minimum of the Morse potential coincide with the scaled WCA by setting *R*_0_ = 2^1*/*6^*σ* + Δ, while the constant *C* shifts the potential to zero at the cutoff *R* = 10*σ*.

### C Noisy Continuous Time Random Walk

In the framework of a nCTRW, the time evolution of the lateral (2D) position of the nanoparticle **r**(*t*) is written as [37, 42]

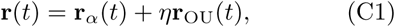

which describes a superposition of a CTRW sub-diffusion process **r**_*α*_(*t*) (with anomalous exponent 0 < *α* < 1) [37, 40, 41] and Ornstein-Uhlenbeck (OU) noise **r**_OU_(*t*) [23, 50] (see fig. 9 for an example trajectory of such a process). The weight *η* > 0 controls the relative contribution of the OU noise to the overall motion. Assuming both of these processes to be independent and letting each start at the origin, we obtain the following expression for the MSD [42]:

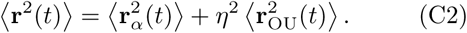

Here brackets denote either an ensemble or a time-ensemble average (which we label with subscripts ‘e’ and ‘te’ respectively), and the averaging for both processes is done with respect to their individual probability density functions (PDFs), i.e. *P*_*α*_(**r**_*α*_, *t*) and *P*_OU_(**r**_OU_, *t*).

The OU process is retrieved by integrating the overdamped Langevin equation [23, 42]

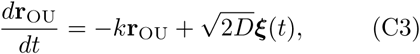

with *k* the stiffness, *D* the diffusion coefficient, and ***ξ***(*t*) = [*ξ*_*x*_(*t*), *ξ*_*y*_(*t*)] a vector of Gaussian white noise processes *ξ*_*i*_(*t*) with zero mean and delta correlation. This equation can be solved for the MSD yielding [23, 42]

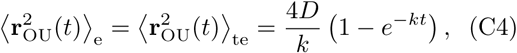

where we have used the ergodicity of the OU process to equate the ensemble average to the time-ensemble average [50].

The CTRW process is described in terms of random jumps Δ**r** and waiting times *t*_*w*_ in between these jumps, which are drawn from independent distributions *p*(Δ**r**) and *ψ*(*t*_*w*_) respectively. In the Fourier-Laplace domain (depicted with the transformation variables **k**, *s*) the PDF of the CTRW relates to the jump and waiting time distributions via the Montroll-Weiss equation [40, 51]

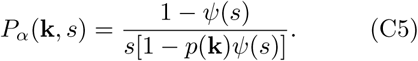

Based on this relation and following the work of [51–53], one can show that for a long-tailed waiting time distribution (asymptotic behavior *ψ*(*t*) ∼ (*t/τ*)^−(1−*α*)^ and *ψ*(*s*) ∼ 1 −(*τs*)^*α*^, with *τ* a decay time scale), the ensemble averaged MSD grows subdiffusively in time according to

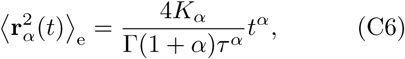

while, due to ergodicity breaking, the time-ensemble average grows linearly in time via

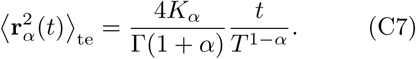

Here *K*_*α*_ is the anomalous diffusion coefficient, Γ the Gamma function, and *T ≫ t* the total time used for time-averaging (the length of individual trajectories).

Combining these results yields an expression for the time-ensemble averaged MSD of the nCTRW:

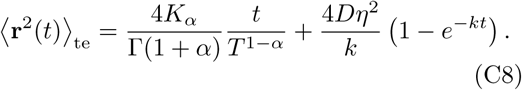

Inspection of this formula reveals three different regimes. Using ⟨ **r**^2^(*t*) ⟩ _te_ ∼ 4*D*_app_*t* we note that for small times *kt* ≪ 1 the MSD grows linearly in time with an apparent diffusion coefficient

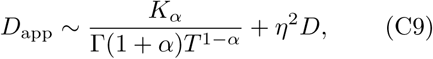

i.e. both the CTRW and OU processes contribute to the MSD. In the long time limit *kt* ≫ 1, the OU contribution to the MSD diminishes resulting in a decreased apparent diffusion coefficient given by

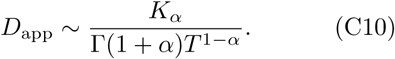

Thus, at intermediate times *kt* ≈ 1 and for strong enough noise *η* there exists a third regime where *D*_app_ decreases in value and as a result the MSD grows subdiffusively in time. Moreover, the expression for the MSD (eq. (C8)) has been confirmed with numerical simulations, indicating the emergence of separate diffusive regimes upon increasing the noise strength (see fig. 9).

### D Supplementary Movies

Supplementary movie 1

Caption: Visualisation of the motion and trajectory of a nanoparticle (colored in magenta) on a fluid membrane consisting of 60% receptors (colored in green). The total time span is on the order of ∼ 12000*τ*.

Supplementary movie 2

Caption: Visualisation of the motion and trajectory of a nanoparticle (colored in magenta) on a fluid membrane consisting of 20% receptors (colored in green). The total time span is on the order of ∼ 12000*τ*.

Supplementary movie 3

Caption: Visualisation of the motion and trajectory of a nanoparticle (colored in magenta) on a gel-like membrane consisting of 20% receptors (colored in green). The total time span is on the order of ∼ 12000*τ*.

